# Metabolically engineered plant cell cultures as biofactories for the production of high-value carotenoid pigments astaxanthin and canthaxanthin

**DOI:** 10.1101/2025.03.18.643922

**Authors:** Bárbara A. Rebelo, M. Rita Ventura, Rita Abranches

## Abstract

Ketocarotenoid pigments, namely astaxanthin and canthaxanthin, are naturally produced by a limited number of species. Demand for these high-value molecules is increasing due to their application in industries such as animal feed, cosmetics, dietary supplements, and pharmaceuticals. We developed a sustainable production platform using metabolically engineered tobacco BY-2 cell suspension cultures by expressing a β-carotene ketolase gene from a marine bacterium and/or overexpress phytoene synthase and desaturase genes. The resulting cell lines exhibited different colors and produced various combinations of ketocarotenoids based on their genetic modifications. Single transformation with the ketolase gene produced salmon-colored cell lines with up to 50 µg g^-1^ DW of canthaxanthin and 127 µg g^-1^ DW of astaxanthin, while expression of all three genes significantly increased canthaxanthin production to 788 µg g^-1^ DW. We demonstrate that undifferentiated cultured plant cells are capable of producing ketocarotenoids, offering an alternative biological solution to natural producers and chemical synthesis.

## Introduction

Plants produce a range of secondary metabolites with numerous applications for humans and animals. Among the various metabolites, carotenoids are of particular importance for living systems due to their antioxidant properties and protective action against photooxidative damage [1,2]. In nutraceuticals and functional foods, consumers seek natural sources of these valuable antioxidants for improved health and wellness [3]. The cosmetics and skincare industries take advantage of carotenoids for their anti-aging and UV-protective properties, meeting growing preferences for natural and sustainable ingredients [4]. Additionally, pharmaceutical companies incorporate carotenoids into formulations for the prevention or treatment of various chronic diseases, including cardiovascular diseases, diabetes, and eye diseases [4–6].

High-value pink-to-red ketocarotenoids, including canthaxanthin and astaxanthin, represent a subgroup of carotenoids widely used in nutraceutical, cosmetic and animal feed industries [7,8]. Ketocarotenoids contain one or more ketone groups, which are functional groups containing a carbonyl group. The characteristic color of ketocarotenoids is determined by their structure. The conjugated double-bond structure acts as a light-absorbing chromophore, while the presence of two keto groups shifts the absorption maximum to higher wavelengths [8].

Canthaxanthin, also known as β,β-carotene-4,4’-dione, results from the modification of β-carotene by the β-carotene ketolase enzyme which adds two oxo substituents at 4- and 4’-positions of the β-ionone backbone. Canthaxanthin is a high-value product with many market applications, but it is also the substrate for obtaining astaxanthin. The enzyme β-carotene hydroxylase catalyzes the introduction of two hydroxyl moieties into the canthaxanthin cyclic rings at 3- and 3’-positions, affording astaxanthin. On the other hand, if β-carotene hydroxylase acts on β-carotene first, it results in the formation of zeaxanthin, that can be modified into astaxanthin by the action of the β-carotene ketolase enzyme (Figure 1). Zeaxanthin undergoes modification by the β-carotene ketolase enzyme with two oxo substituents added at 4- and 4’-positions of the β-ionone backbone [8–10]. In the case of astaxanthin, the presence of two additional alpha hydroxy groups may contribute to its enhanced antioxidant capacity [8,10].

**Figure 1.**
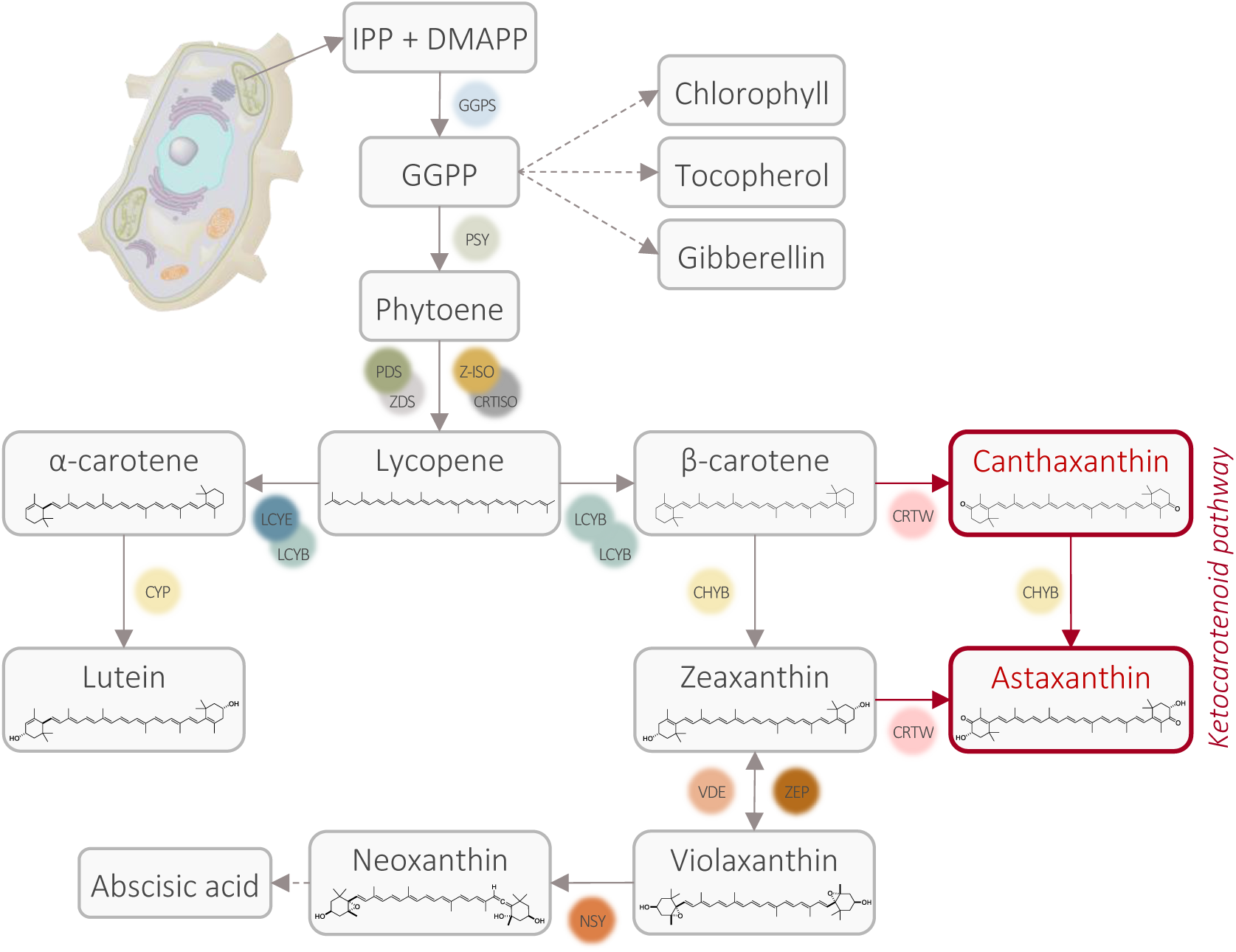
Schematic overview of the carotenoid biosynthetic pathway in plants. The isopentenyl diphosphate (IPP) and dimethylallyl diphosphate (DMAPP) building blocks can be produced from the methylerythritol phosphate (MEP) cytoplasmic and mevalonate (MVA) plastid pathways. The grey boxes indicate the main carotenoids and derivatives found in plant systems, while the red boxes indicate the heterologous ketocarotenoid pathway. Abbreviations of enzymes: GGPS (geranylgeranyl pyrophosphate synthase), PSY (phytoene synthase, CRTB for bacteria), PDS (phytoene desaturase, CRTI for bacteria), ZDS (ζ-carotene desaturase, CRTI for bacteria), Z-ISO (ζ-carotene isomerase, CRTI for bacteria), CRTISO (carotenoid isomerase, CRTI for bacteria), LCYE (lycopene ε-cyclase), LCYB (lycopene β-cyclase, CRTY for bacteria), CYP (cytochrome P450 enzyme), CHYB (β-carotene hydroxylase, BHY for algae, CRTZ for bacteria), CRTW (β-carotene ketolase, BKT for algae), VDE (violaxanthin de-epoxidase), ZEP (zeaxanthin epoxidase), NSY (neoxanthin synthase). This visual representation shows the stepwise process involved in ketocarotenoid synthesis.

Only a few species of algae and microorganisms are able to perform the ketolation reaction necessary to produce these ketocarotenoids [8]. Examples of natural producers include the microalgae *Haematococcus pluvialis* and *Chromochloris zofingiensis*, the bacteria *Brevundimonas* sp. SD212, and the yeast *Phaffia rhodozyma*. The only known plant genus naturally producing ketocarotenoids is *Adonis*, which accumulates high levels of astaxanthin in its flower petals [11]. In contrast to the typical pathway involving β-carotene hydroxylase and ketolase, this plant utilizes the β-ring 4-dehydrogenase (CBFD) and 4-hydroxy-β-ring 4-dehydrogenase (HBFD) enzymes. Nonetheless, natural sources of ketocarotenoids have been overlooked as major producers due to the high production costs, insufficient quantities produced by these organisms, and the complexity of isolating ketocarotenoids [10]. For example, *H. pluvialis* cultivation for astaxanthin production requires high light, controlled nutrient depletion, and extended growth periods, resulting in limited output and consequently a higher price for natural astaxanthin [12].

Thus, the feed industry often resorts to chemical synthesis for their supply [8,13], raising sustainability concerns due to its reliance on petrochemical-derived precursors. Moreover, the structural complexity of ketocarotenoids adds to the laborious nature of their production by chemical synthesis, as it requires long synthetic routes and highly stereoselective reactions, making large-scale production challenging and costly [14–16]. Nevertheless, biological production of ketocarotenoids cannot economically compete with synthetic production.

Various strategies have been employed to improve ketocarotenoid production with the goal of optimizing resource utilization and minimizing adverse effects. Strategies of genetic modification have been accomplished, affording hydroxyl/ketolated carotenoids with variable qualitative and quantitative profiles (reviewed in detail in [1,8,9]). A noteworthy example is the orange-red-grained astaxanthin rice [17], obtained through transformation with multiple carotenogenic genes: plant phytoene synthase (*psy*), bacterial phytoene desaturase (*crtI*) and algal β-carotene hydroxylase (*bhy*) and ketolase (*bkt*). This approach generated various germplasms capable of producing astaxanthin (Astx = 16 µg g^-1^ DW), although the expression of endogenous genes influenced the final outcome [17]. In another study, Mortimer and co-workers introduced bacterial β-carotene hydroxylase and a ketolase gene (*crtZ* and *crtW*) from *Brevundimonas* sp. into *Nicotiana glauca,* resulting in astaxanthin production of 0.140 mg g^-1^ DW [18]. Using the same enzymes in tomato, Enfissi and colleagues achieved an even higher astaxanthin yield of 0.362 mg g^-1^ DW [19]. Moreover, Nogueira and colleagues improved the ketocarotenoid content in tomato by crossing a low ketocarotenoid producing line (ZW) [19] with an orange fruit recombinant inbred line that accumulated high levels of β-carotene [16,20]. More recently, Tanwar and colleagues [21] reported ketocarotenoid production in chloroplast-engineered tobacco plants expressing a *bkt* gene in combination with the lycopene cyclase (*lcy*) and *bhy* genes. However, they obtained higher xanthophyll levels rather than ketocarotenoids, and only reported the presence of 4-ketolutein. They reasoned that 4-ketolutein was obtained probably due to the presence of either α-carotene or lutein in the plant system, and that the β-carotene ketolase enzyme would show a preference to oxidize only one ring over two rings present in zeaxanthin. Using a different approach, Allen and co-workers engineered *Nicotiana benthamiana* plants with β-ring 4-dehydrogenase (CBFD) and 4-hydroxy-β-ring 4-dehydrogenase (HBFD) enzymes from *Adonis aestivalis*, in order to produce astaxanthin directly from β-carotene [22]. Drawbacks in carotenoid expression have been reported, such as growth reduction in transformed *Brassica napus* and a decline in chlorophyll content in the leaves, without causing a significant impact on photosynthetic efficiency [23,24].

Apart from plants, microalgae and fermentation systems, including bacteria and yeast, have also been used. Chen and colleagues observed an enhanced astaxanthin production in *Chlamydomonas reinhardtii* by overexpressing either endogenous or *Phaffia rhodozyma* β-carotene ketolase genes, observing greater effects with the heterologous enzyme [25]. In the same microalgal model, combining *C. reinhardtii* β-carotene *bkt* with phytoene synthase (*crtB*) from *Pantoea ananatis*, and *C. reinhardtii* β-carotene 3-hydroxylase (*chyb*) [26] resulted in a volumetric astaxanthin production of 9.5 mg L^-1^ (4.5 mg g^-1^ cell dry weight (DW)) under mixotrophic conditions and 23.5 mg L^-1^ (1.09 mg L^-1^ h^-1^) under high cell density conditions. Regarding intracellular storage, in nonoleaginous organisms such as *Saccharomyces cerevisiae*, the number and size of lipid droplets are limited due to the low lipid content, and the heterologous lipophilic products are stored primarily in cell membranes, which can place a heavy metabolic burden on the chassis cells [27].

Recent advancements in plant synthetic biology provide abundant genetic materials and engineering strategies, creating opportunities for tailored cell factories for specific applications [28]. Plant-based systems for carotenoid production offer environmentally friendly alternatives to synthetic production methods, aligning with the increasing consumer preference for natural and sustainable products. These systems can be scaled up, enhancing economic viability by optimizing production processes, reducing costs, and ensuring a reliable supply of carotenoids from renewable plant sources. This approach supports a more sustainable and ecologically responsible method of meeting market demands while reducing the ecological footprint of carotenoid production. In the present study, we conducted metabolic engineering to produce ketocarotenoids in plant cell suspension cultures, namely tobacco BY-2 cells. Carotenoid production, without substrate feeding, was achieved by expressing various combinations of carotenoid-related genes. This combinatorial transformation resulted in colored BY-2 cells that exhibited continuous pigment production throughout subculturing, while maintaining normal cell growth and developmental process. By using this platform, it is possible to provide ketocarotenoid production in a reliable, reproducible, efficient, with rentable yields, whilst maintaining an environmentally friendly sustainable process that addresses the limitations of other systems.

## Results

### Vector construction for the expression of selected genes involved in carotenoid biosynthesis

The non-green BY-2 cell line, derived from *Nicotiana tabacum* plants, is widely used as an undifferentiated cell model for many fundamental biology studies. However, there are only a few reports on the heterologous biosynthesis of pigments in these cultured cells [29–31]. In this work, we expressed key carotenogenic genes to complete and enhance the carotenoid pathway for the production of astaxanthin and canthaxanthin. We selected the β-carotene ketolase gene (*crtW*) from the marine bacteria *Brevundimonas* sp. strain SD212. The introduction of this gene was essential for ketocarotenoid biosynthesis, as BY-2 cells do not naturally possess a β-carotene ketolase gene in their genome. We also analyzed endogenous carotenogenic transcripts from BY- 2 cells and compared the results with the literature. Since previous studies have shown that boosting substrate production upstream of the ketocarotenoid pathway can enhance carotenoid biosynthesis yields, we reasoned that co-expressing transgenes for phytoene synthase and desaturase would increase the availability of phytoene and lycopene. Therefore, we selected the phytoene desaturase gene (*crtI*) from *Pantoea ananatis* and the phytoene synthase gene (*psy*) from *Zea mays*. The *psy* or *crtI* genes were inserted into the binary vector pK2GW7, while the synthetic *crtW* gene was cloned into the pTRA vector. This resulted in constructs pY, pI and pW (Figure 2), with all genes under the control of the 35S promoter from cauliflower mosaic virus. The constructs were named based on the last letter of each gene included in the cassette. To direct the enzyme to the plastids, a plastid-transit peptide from the pea Rubisco small subunit was fused to the non-plant derived *crtI* and *crtW*, while *psy* already contained an endogenous transit peptide.

**Figure 2.**
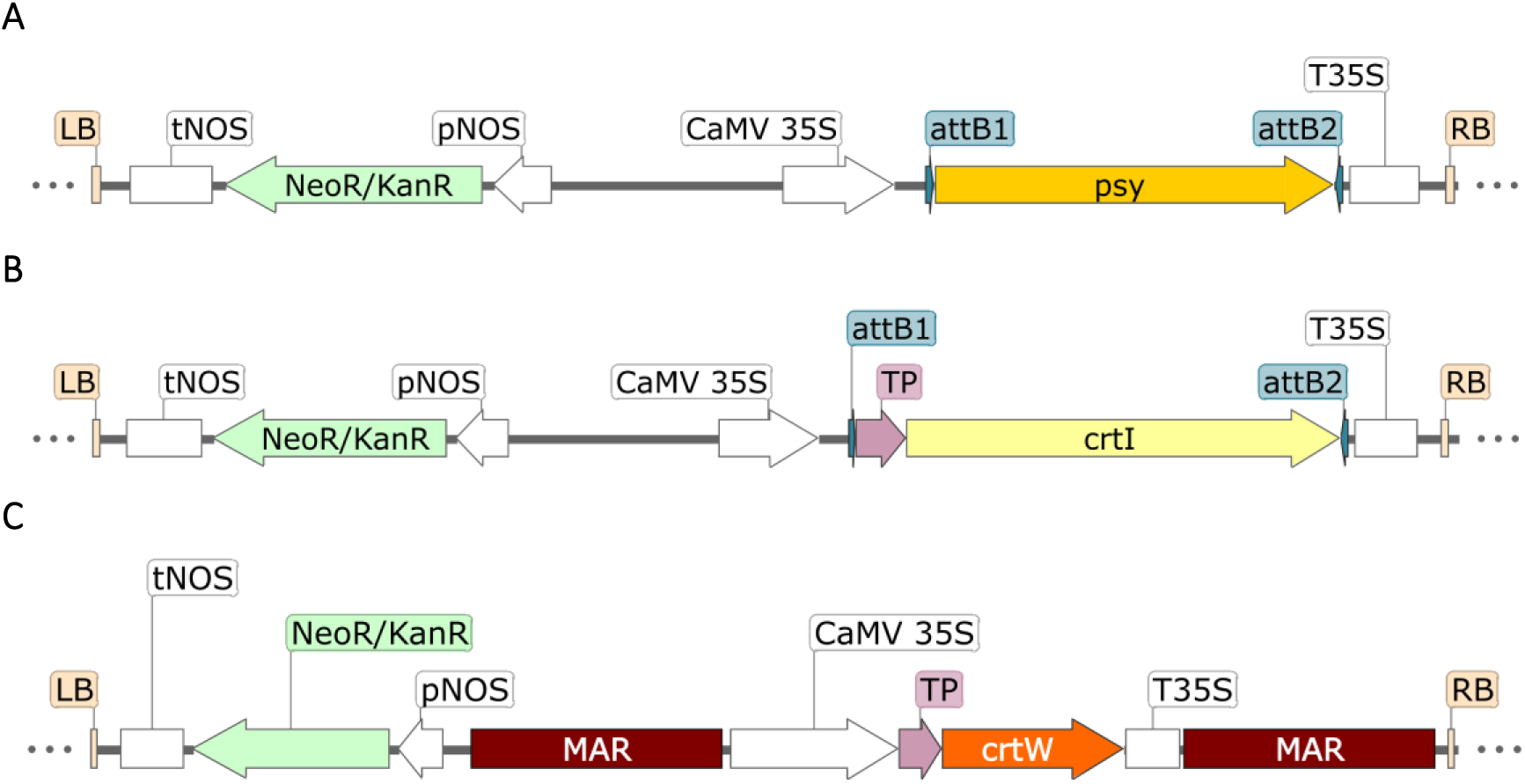
Schematic representation of the T-DNA cassette of (A) construct pY, (B) construct pI and (C) construct pW used for the transformation of BY-2 cells. Regions are represented as follows: left border (LB), nopaline synthase terminator (tNOS), kanamycin resistance marker (NeoR/KanR), nopaline synthase promoter (pNOS), matrix attachment regions (MAR), Cauliflower Mosaic Virus 35S promoter (CaMV 35S), site-specific recombination site (attB1 and attB2), transit peptide (TP), phytoene synthase coding sequence (AY324431), phytoene desaturase coding sequence (D90087), β-carotene ketolase coding sequence, 35S terminator (T35S), right border (RB). All dark lines are nucleotide sequences without relevant features.

### *Nicotiana benthamiana* allowed for the rapid functional analysis of carotenogenic genes involved in ketocarotenoid biosynthesis

*Nicotiana benthamiana* is a well-established biofactory system for transient expression, characterized by large leaves that facilitate infiltration and rapid production of recombinant proteins. This species was selected due to its natural ability to supply the precursor geranylgeranyl pyrophosphate (GGPP) and carotenoids, which support the synthesis of ketocarotenoids. In order to assess the effect of expressing the heterologous genes *psy*, *crtI* and *crtW*, we performed transient expression experiments using Agrobacteria-mediated delivery in fully developed leaves. Since mature chloroplasts in these leaves naturally synthesize carotenes and xanthophylls, we aimed to determine if the introduced genes could extend the pathway and produce ketocarotenoids, thereby altering the carotenoid composition. Furthermore, we investigated whether endogenous enzymes, in conjunction with the β-carotene ketolase would be sufficient for ketocarotenoid production. We tested single-gene constructs (pY; pI; pW) and multi-combinatorial (pY + pI; pY + pW; pI + pW; pY + pI + pW).

High-performance liquid chromatography (HPLC) analysis of ethanolic leaf extracts was performed to qualitatively evaluate the carotenoid profile, with each leaf serving as an independent biological replicate. Based on retention time (Figure S1), absorption spectra, and comparison with available standards, smaller amounts of neoxanthin (Neo), violaxanthin (Vio) and β-carotene (β-car) were identified. Additionally, we analyzed the pigment profile of Arabidopsis leaves and *Tetraselmis chuii* total carotenoid extract using the same HPLC conditions, comparing them with the carotenoid profiles described in the literature. Using this data, along with information on the different polarities of the pigment molecules and available carotenoid descriptions in literature reports (e.g. [17,23,32]), we identified putative zeaxanthin/lutein (*Z/L), chlorophyll b (*Chl b) and chlorophyll a (*Chl a). Interestingly, trace amounts of two carotenoids (peak 1 and 2, Figure S1) were detected exclusively in the leaves infiltrated with the *crtW* gene, suggesting ketolase-mediated modification of carotenoid molecules. While the carotenoid corresponding to peak 3 (Figure S1) was detected in other samples, it accumulated more in leaves expressing *crtW*. Free astaxanthin was not identified in the analyzed extracts. Peaks 1 and 2 may represent ketocarotenoid intermediates since astaxanthin biosynthesis requires four enzymatic steps from β-carotene, divided by two different pathways. These peaks might also correspond to astaxanthin derivatives with different chemical groups attached to their β-rings hydroxyl groups. Astaxanthin can be extracted in the free form, which is more polar, or as fatty acid mono- and di-esters [33], which is less polar and exhibiting higher retention times. With these results, we proceeded with the transformation of plant cell cultures using these constructs.

### Elicitation experiments of carotenoid synthesis in undifferentiated non-photosynthetic BY-2 cells

To investigate the effects of *crtW* in non-green tissues characterized by rapid growth and controlled conditions, tobacco BY-2 cultured cells were selected as a potential platform for ketocarotenoid production. Previous studies have shown that BY-2 cells grown in darkness do not accumulate specialized carotenoids [34,35], including key precursors for ketocarotenoids biosynthesis, despite the endogenous expression of phytoene synthase (*psy*) and desaturase (*pds*), lycopene β-cyclase (*lcyb*), and β-carotene hydroxylase (*chyb*) transcripts. Since carotenoids play a role in photoprotection, light exposure was hypothesized to induce carotenoid biosynthesis. Consequently, BY-2 cells were transferred into liquid MS medium with reduced sucrose (2% (w/v)) and grown under controlled light conditions. Cell suspension cultures were refreshed by transferring 3% (v/v) of cells into 25 ml of MS medium, with a stepwise 50% reduction in sucrose content (Table S1), following the methodology outlined in [36]. This approach, previously applied to Arabidopsis, promoted the differentiation of chloroplasts from proplastids. We also tested various kinetin concentrations (0 to 27.88 µM), since this artificial cytokinin is known to aid proplastid differentiation [37]. We aimed to identify optimal conditions for enhancing carotenoid biosynthesis while maintaining cell viability. Following a preliminary analysis of BY-2 cell viability and total carotenoid content, specific samples were collected and subjected to HPLC analysis to assess the new carotenoid profile in BY-2 cells.

After four months, carotenoid production was observed under optimal growth conditions. HPLC analysis of specific samples (Figure S2) revealed the presence of at least three carotenoids, identified as neoxanthin, violaxanthin and β-carotene, based on spectral properties and co-chromatography with standards. These were detected in cultures grown with 3% sucrose and 9.29 µM of kinetin, 1% sucrose and 13.94 µM of kinetin, and 0.5% sucrose and 18.59 µM of kinetin. As high kinetin concentrations and lower sucrose concentrations reduced cell division, we lowered kinetin concentrations for better cell viability. After approximately one year, BY-2 cells grown under a photoperiod exhibited the capability to produce carotenes and xanthophylls (Figure S2), which were absent in the WT cells grown in the dark. Moreover, chlorophyll was not detected throughout the experiment.

### Combinatorial nuclear transformation of BY-2 cells generated distinct phenotypes

The carotenoid pathway of BY-2 cells was metabolically engineered through combinatorial transformation with carotenogenic genes, all under the control of the strong promoter CaMV 35S. Seven independent transformation events were conducted (Table S2), involving the individual transformation of BY-2 cultured cells with phytoene synthase (*psy*), phytoene desaturase (*crtI*), or β-carotene ketolase (*crtW*) genes from *Zea mays*, *Pantoea ananatis*, and *Brevundimonas* sp. SD212, respectively. Combinatorial transformations involved two genes in the following combinations: *psy* with *crtI*, *psy* with *crtW*, and *crtI* with *crtW*. A final multigene transformation included all three genes. For the identification of these cell lines, we used the last letter of each gene introduced to name the cell line, as follows: "Y" represents cells with the heterologous *psy* gene, "I" represents *crtI*, and "W" represents *crtW*. When two or more genes were combined, the letters were combined accordingly: "YI" refers to cell lines with both *psy* and *crtI*, "YW" refers to cells with *psy* and crtW, "IW" refers to cell lines with both *crtI* and *psy*, and "YIW" designates cells containing all three genes, *psy*, *crtI*, and *crtW*. These names (Y, I, W, YI, YW, IW, and YIW) will be used throughout the text to refer to each specific transformation event (Figure 3A). The numbers following these letters indicate the specific *calli* selected. To target the bacterial CRTI and CRTW enzymes to the plastids, a transit peptide from the pea RuBisco small subunit was included in the expression cassettes. This transit peptide has been successfully used in various systems, including in rice endosperm, maize, and soybean [17,32,38]. The maize *psy* gene contains an endogenous transit peptide and was used without modifications. Numerous *calli* were generated from each transformation and underwent several rounds of antibiotic selection before the screening.

**Figure 3.**
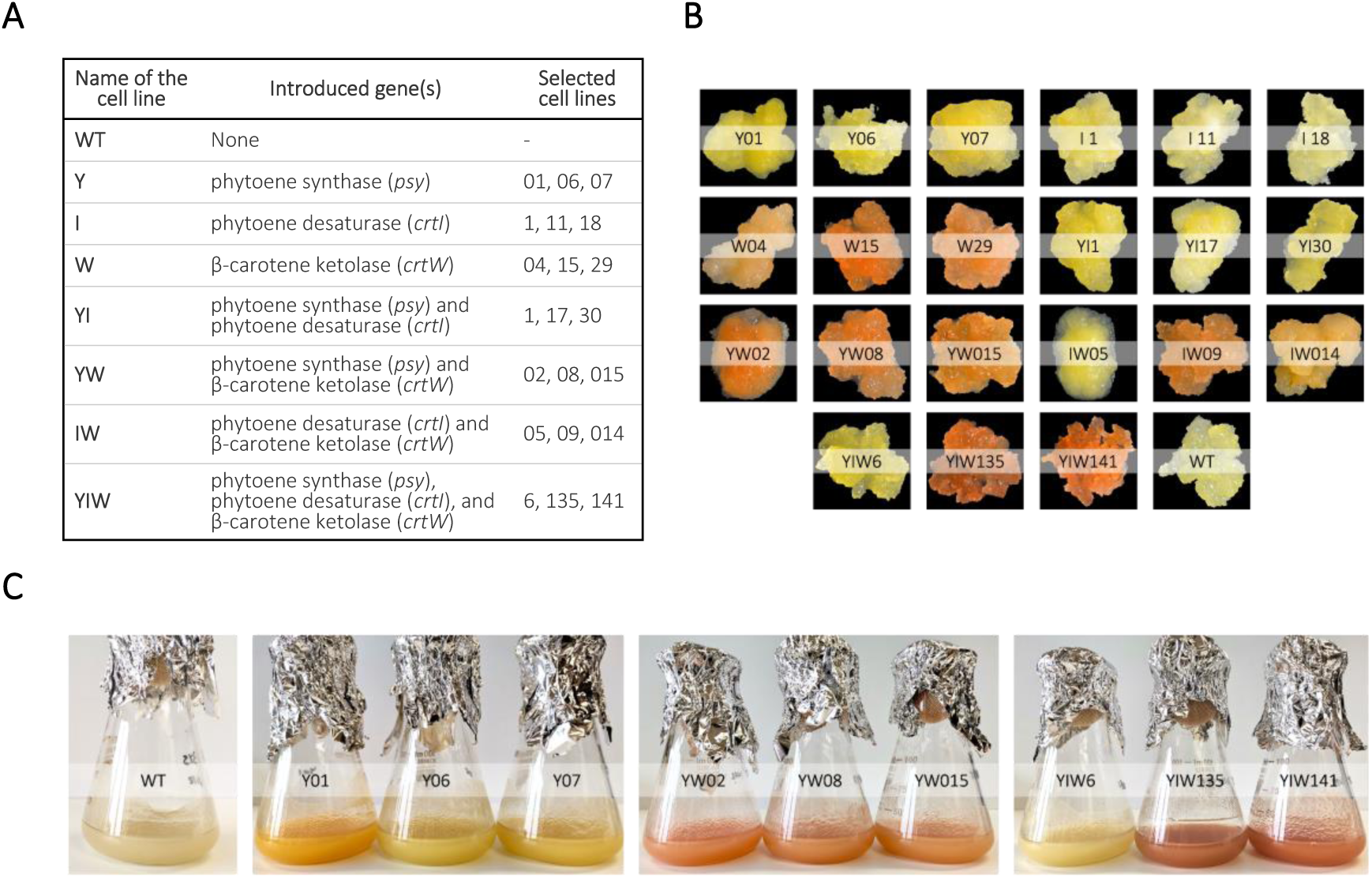
Phenotype of *Nicotiana tabacum* BY-2 cell lines compared to the wild-type. (A) Classification of BY-2 cell lines containing the heterologous *psy*, *crtI*, and/or *crtW* genes resulting from the single and combinatorial transformations with the pY, pI, and/or pW vectors. (B) Pale, yellow or pink-orange pigmented tobacco BY-2 *calli*. (C) BY-2 cultured cells, the WT and the highest yielding cell lines for the production of xanthophylls, astaxanthin, and canthaxanthin.

The newly established transgenic BY-2 lines showed distinct phenotypes among the different transformation events and compared to pale-yellow wild-type *calli*. *Calli* transformed with *crtI* resembled the wild-type, whereas those transformed with *psy* displayed light to dark yellow pigmentation. In contrast, *calli* transformed with the *crtW* gene exhibited light brownish, salmon, or orange coloration (Figure 3B). Co-expression of two or three genes produced various colorations: white or yellow (transformation with constructs pY + pI), white or pink-orange (transformation with constructs pY + pW or with pI + pW) and white, yellow or pink-orange colored *calli* (transformation with constructs pY + pI + pW). Despite these color changes, all the transgenic BY-2 lines exhibited normal growth and morphology similar to wild-type *calli*. The intensity of pigmentation in transgenic *calli* increased after three weeks of growth under controlled light conditions. Several *calli* from each transformation were selected based on color. No modifications to the culture medium, such as adjustments in sucrose or kinetin concentrations, were made, as the *calli* exhibit color upon light exposure post-transformation. However, kinetin potential as an elicitor to enhance pigment production could be explored in future studies.

The presence of transgenes was confirmed by PCR using specific primers, and no bands corresponding to *psy*, *crtI* or *crtW* were detected in the wild-type (negative control) (Figure S3). A total of twenty-one independent lines were selected, and liquid cultures were established (Figure 3C, Figure S4). Importantly, expression of the heterologous genes did not affect growth rate or biomass production compared to WT cell suspension culture. Both WT and transgenic BY-2 *calli* were subcultured every three weeks, and cell suspensions were subcultured on the 7^th^ day of the growth curve.

### Engineering of the carotenoid biosynthetic pathway led to the production and accumulation of ketocarotenoids

To characterize the carotenoid profiles of WT and transgenic BY-2 cell lines, cells were harvested on the 7^th^ day of the growth curve. Pigment analysis was performed at this phase, because a progressive increase in color intensity was observed throughout the growth period. HPLC analysis revealed a correlation between the observed coloration and the resultant carotenoid profile. In the WT carotenoid extract, three main pigments were detected: the xanthophylls neoxanthin and violaxanthin, alongside β-carotene (Figure 4, Figure S5). In previous work, chlorophylls and these xanthophylls were identified on green microalgae, tobacco and Arabidopsis leaves (results not shown). By leveraging all the collected information, it was possible to qualitatively identify the two xanthophylls without relying on standards. As expected, no astaxanthin or canthaxanthin were detected in the WT extracts.

**Figure 4.**
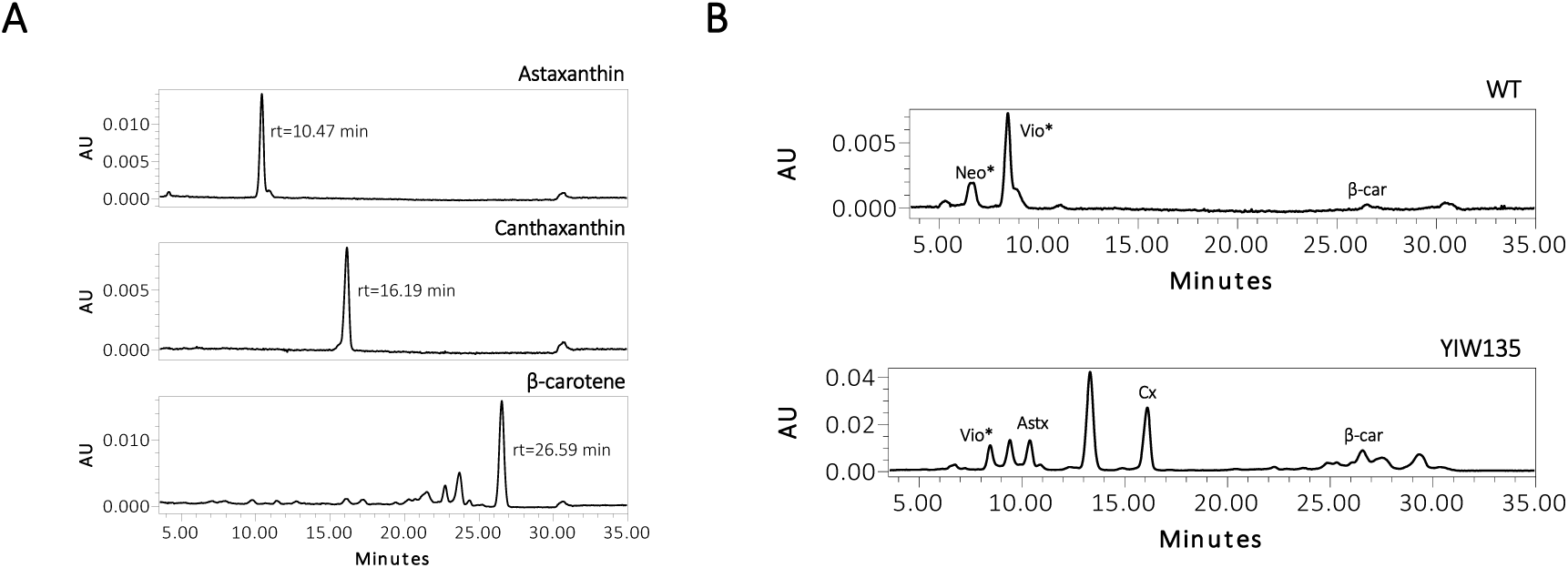
Carotenoid profile of tobacco BY-2 wild-type (WT) and the highest ketocarotenoid-producing line (YIW135). The resulting chromatograms were recorded at a wavelength of 445 nm, with the y-axis scaled to the highest peak to enhance clarity. The carotenoids identified were putative neoxanthin (*Neo), putative violaxanthin (*Vio), astaxanthin (Astx), canthaxanthin (Cx) and β-carotene (β-car).

The tobacco lines I, Y and YI exhibited similar pigment compositions, featuring putative neoxanthin, violaxanthin and β-carotene as the predominant pigments. Although these lines shared a similar carotenoid profile, their colors varied due to differences in the accumulation of each pigment. The cell lines Y displayed a bright lemon-yellow color, likely reflecting a higher accumulation of xanthophylls, while the cell lines YI showed a darker yellow coloration. Moreover, these lines exhibited an increased number of compounds eluting between 17 and 25 minutes in contrast to the cell lines I. These compounds could potentially be violaxanthin fatty acid esters, which were also identified in *Nicotiana glauca* leaves [23]. The xanthophylls zeaxanthin and lutein, isomers present in *N. tabacum* leaves [39], are challenging to distinguish due to a structural difference involving a double bond within one of the ionone rings (ε-ring). Co-chromatography with a lutein standard confirmed its absence in tobacco BY-2 lines. Although HPLC analysis with a zeaxanthin standard was not performed, the synthesis of zeaxanthin in BY-2 cells is supported by the presence of its precursor, β-carotene, and its derivative, violaxanthin.

The introduction of the β-carotene ketolase enzyme (CRTW) completed the carotenoid biosynthetic pathway, resulting in the production of ketocarotenoids. Cell lines W, expressing the *crtW* gene, exhibited distinct carotenoid profiles compared to the wild-type, and all produced both astaxanthin and canthaxanthin, with several additional compounds detected. For the HPLC analysis, the total carotenoid extract was separated using a C18 column, which is a reverse phase column with a non-polar stationary phase. Due to the elution conditions employed in this methodology, more polar compounds (such as xanthophylls) eluted earlier than non-polar compounds (carotenes). We hypothesize that the compounds with a retention time between 9 and 15 min could be intermediaries in the ketocarotenoid pathway, such as adonirubin or adonixanthin. This assumption is based on the polarity of the molecules, existing literature [40–42], and the understanding that not all substrates are converted into the final product of the pathway (astaxanthin).

Quantitative data showing the levels of astaxanthin, canthaxanthin and β-carotene produced by wild-type and transgenic cell cultures is summarized in Table 1. A notable increase in β-carotene levels was observed in cell lines expressing *psy*, either singly or in combination with *crtI* gene. In the transgenic lines Y and YI, β-carotene levels increased 20- to 60-fold compared to wild-type cells. In the cell lines YIW, co-expressing all three genes resulted in an increase in β-carotene up to 55-fold. Transformation with the *crtW* gene alone enabled BY-2 cells to produce ketocarotenoids (Figure S5), using naturally synthesized xanthophylls as substrates. The astaxanthin content in the cell lines W ranged from 90 to 127 µg g^-1^ DW, while canthaxanthin amounts ranged from 20 to 55 µg g^-1^ DW (Table 1, Figure 5). The optimal combination for astaxanthin production was achieved through the co-expression of *crtW* with *psy*. In comparison to cell line W04, YW02 exhibited a 1.9-fold increase in astaxanthin and a 2.4-fold increase in canthaxanthin accumulation. Conversely, co-expression of *crtW* with *crtI* (lines IW) resulted in decreased ketocarotenoid levels. Notably, co-expression of all three genes (lines YIW) led to a significant increase in canthaxanthin production. Specifically, YIW135 and YIW141 accumulated 787.9 and 279.0 µg g^-1^ DW of canthaxanthin, representing a 39-fold and 14-fold increase, respectively, compared to W04.

**Figure 5.**
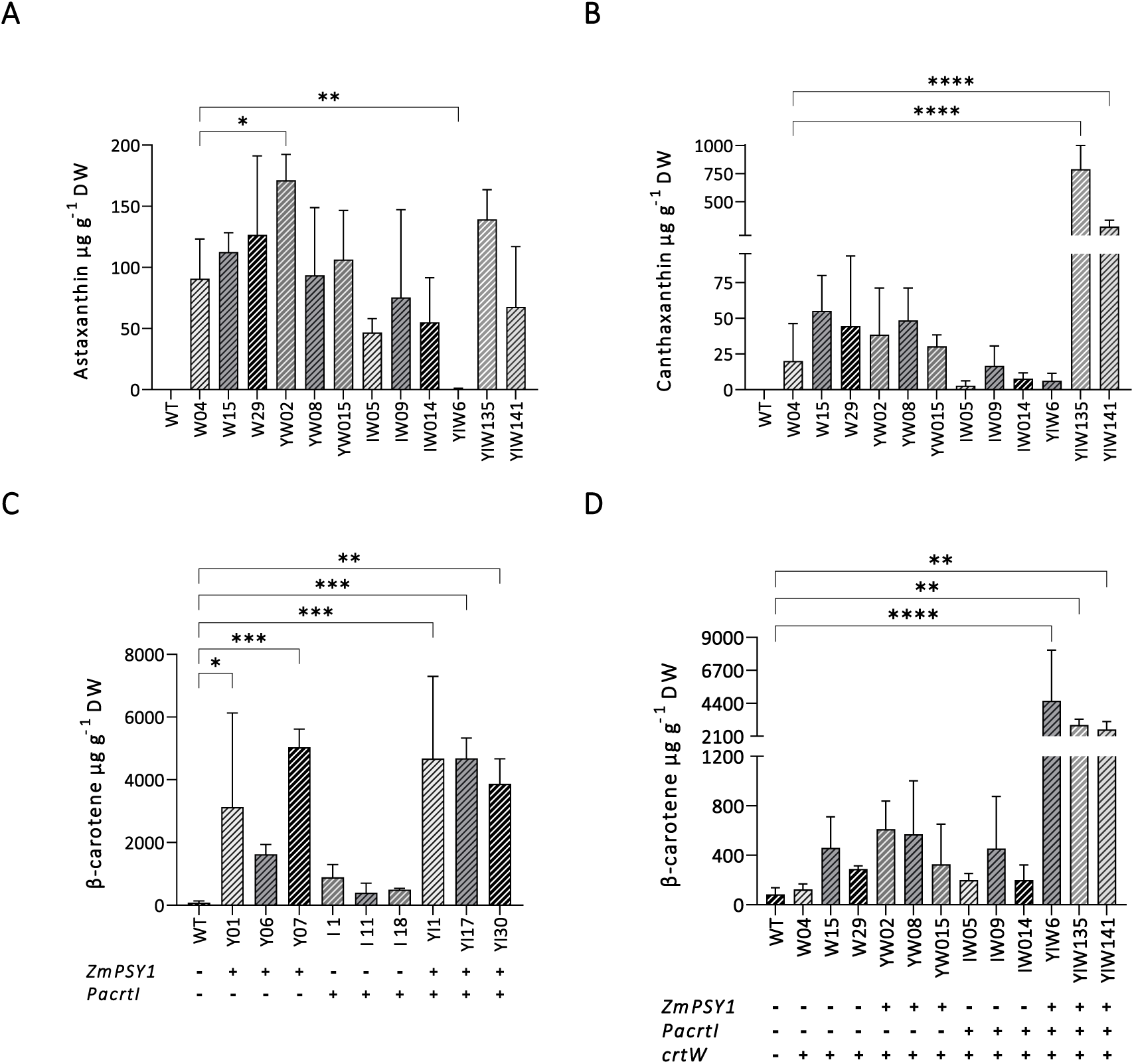
Carotenoid content in tobacco BY-2 cell lines is shown as µg g^-1^ DW (micrograms per gram of dry weight). Quantification of (A) astaxanthin and (B) canthaxanthin in ketocarotenoid producing cell lines. Quantification of β-carotene in (C) xanthophyll and (D) ketocarotenoid producing cell lines. The presence (+) or absence (-) of the heterologous genes is indicated for each line. Each data point represents the mean result derived from three biological replicates from the HPLC run, with error bars indicating the standard deviation (SD) for each measurement. Data exhibiting significant differences are indicated by the asterisk symbol. Statistical analysis was conducted by comparing the data to the cell line W04. The exception to this was the β-carotene quantification in the xanthophyll lines (C), where the statistical analysis was conducted by comparing the data to the WT. The P-Values, as determined by Uncorrected Fisher’s LSD, are indicated as follows: *0.05, **0.01, ***0.0005 and ****<0.0001.

**Table 1.** Carotenoid content in wild-type and three independent transgenic cell lines for each transformation event. Carotenoid levels are represented as µg g^-1^ DW and expressed as a percentage of total carotenoid extract (% TCE)^a^. Each value in carotenoids is the mean result from three biological replicates ± SD. The P-Values, as determined by Uncorrected Fisher’s LSD, are indicated as follows: *0.05, **0.01, ***0.0005 and ****<0.0001. nd: not detected.

| | Astaxanthin | | Canthaxanthin | | β-carotene | |
| --- | --- | --- | --- | --- | --- | --- |
| | $\mu\text{g g}^{-1}$ DW | % of TCE | $\mu\text{g g}^{-1}$ DW | % of TCE | $\mu\text{g g}^{-1}$ DW | % of TCE |
| **Wild-type** | nd | - | nd | - | 83.6 ± 53.4 | 2.6 ± 0.9 |
| **Y01** | nd | - | nd | - | 3 128.5* ± 3 001.2 | 6.0 ± 5.2 |
| **Y06** | nd | - | nd | - | 1 624.2 ± 315.5 | 20.7 ± 4.2 |
| **Y07** | nd | - | nd | - | 5 035.8*** ± 578.8 | 26.0 ± 2.5 |
| **I 1** | nd | - | nd | - | 891.6 ± 408.1 | 12.1 ± 2.7 |
| **I 11** | nd | - | nd | - | 395.7 ± 309.7 | 12.6 ± 8.0 |
| **I 18** | nd | - | nd | - | 496.1 ± 42.1 | 10.9 ± 0.9 |
| **W04** | 90.8 ± 32.4 | 15.5 ± 2.2 | 20.0 ± 26.3 | 1.4 ± 1.2 | 124.2 ± 45.2 | 1.0 ± 0.4 |
| **W15** | 112.6 ± 15.8 | 16.3 ± 2.5 | 55.1 ± 24.7 | 2.8 ± 1.3 | 460.9 ± 249.5 | 2.9 ± 1.3 |
| **W29** | 126.5 ± 64.8 | 22.9 ± 3.2 | 44.5 ± 49.1 | 2.5 ± 1.6 | 289.2 ± 25.0 | 2.6 ± 1.5 |
| **YI1** | nd | - | nd | - | 4 675.9*** ± 2 624.0 | 33.4 ± 1.9 |
| **YI17** | nd | - | nd | - | 4 685.5*** ± 642.2 | 36.3 ± 1.5 |
| **YI30** | nd | - | nd | - | 3 872.9** ± 797.3 | 30.1 ± 3.7 |
| **YW02** | 171.3* ± 21.2 | 14.8 ± 3.6 | 38.4 ± 32.7 | 1.6 ± 1.7 | 611.5 ± 227.9 | 3.0 ± 1.1 |
| **YW08** | 93.5 ± 55.5 | 11.3 ± 0.4 | 48.5 ± 22.7 | 2.1 ± 0.2 | 570.6 ± 431.7 | 3.1 ± 2.0 |
| **YW015** | 106.3 ± 40.3 | 15.2 ± 1.4 | 30.4 ± 7.9 | 1.7 ± 0.5 | 326.8 ± 323.9 | 1.8 ± 1.2 |
| **IW05** | 46.7 ± 11.4 | 11.9 ± 1.0 | 2.8 ± 3.3 | 0.4 ± 0.3 | 199.7 ± 53.3 | 2.1 ± 1.1 |
| **IW09** | 75.4 ± 71.9 | 12.2 ± 2.3 | 16.7 ± 13.9 | 1.3 ± 0.6 | 455.1 ± 421.9 | 2.1 ± 1.2 |
| **IW014** | 55.0 ± 36.6 | 19.6 ± 0.6 | 7.8 ± 4.1 | 1.3 ± 0.1 | 199.0 ± 122.1 | 1.7 ± 1.2 |
| **YIW6** | 0.5** ± 0.8 | 0.2 ± 0.4 | 6.2 ± 5.3 | 0.3 ± 0.3 | 4 584.0**** ± 3 517.8 | 31.9 ± 7.1 |
| **YIW135** | 139.3 ± 24.3 | 7.1 ± 0.3 | 787.9**** ± 213.3 | 12.9 ± 2.3 | 2 878.4** ± 413.8 | 8.0 ± 1.3 |
| **YIW141** | 67.8 ± 49.3 | 5.4 ± 0.2 | 279.0**** ± 56.9 | 5.9 ± 0.7 | 2 578.9** ± 551.9 | 9.5 ± 0.3 |

<sup>a</sup> The sum of the integrated area of each peak corresponds to the total carotenoid content for each sample. To determine the percentage of each specific carotenoid (astaxanthin, canthaxanthin, or β-carotene) within the total extract, the area of each peak was divided by the total peak area.

Among the population of transgenic BY-2 lines examined, one line stood out. Cell culture YIW6 displayed a yellow color. PCR analysis of the genomic DNA confirmed the presence of the *crtW* gene, which was further validated by HPLC. However, both carotenoid profile and quantitative data indicated that β-carotene and zeaxanthin were not efficiently converted into ketocarotenoids, resulting in an accumulation of β-carotene. The amounts of astaxanthin and canthaxanthin detected in YIW6 were merely 0.5 and 6.2 µg g^-1^ DW, respectively. Further studies are necessary to investigate whether the variability in ketocarotenoid accumulation is attributed to factors such as transgene copy number, insertion site, transcriptional-level silencing, or enzyme efficiency.

## Discussion

The growing market demand for carotenoids in various industries reflects a consumer preference for natural, health-promoting compounds. Overall, there is a recognized need to improve and increase carotenoid production, both from natural sources or via heterologous systems, to overcome the drawbacks of natural production, such as high production costs and the complexity of isolating ketocarotenoids. Alternative biological production systems are being explored to achieve cost-effective ketocarotenoid synthesis, prioritizing environmental friendliness and safety for human use. In this context, several efforts have been explored to improve natural large-scale carotenoid production, including overexpressing rate-limiting enzymes, downregulating competing pathways, and enhancing carotenoid storage capacity [43].

### Unveiling ketocarotenoid production in non-photosynthetic undifferentiated cells

Tobacco BY-2 cell suspension cultures are well-established platforms for molecular farming, particularly for producing recombinant proteins. This study proposes that these cultured cells hold potential for producing secondary metabolites, in particular carotenoid pigments. Compared to other expression systems, BY-2 cells offer advantages such as fast growth, controlled and confined environment, reduced costs, adherence to good manufacturing practices (GMP) and safety [44,45]. Typically, tobacco BY-2 cells grow in the dark, in MS medium supplemented with a carbon source, vitamins and auxins. As non-photosynthetic cells, they do not naturally produce and store specific secondary metabolites intricately linked with photosynthesis, like specialized carotenoids, which are normally involved in photoprotection. In higher plants, light is known to influence the regulation of carotenoid biosynthesis and the expression of carotenogenic genes [46]. In this study, tobacco BY-2 cell cultures were transitioned from total darkness to a 16-hour light period, resulting in the production of carotenes and xanthophylls. The localization of carotenoids in BY-2 cells is under investigation, likely plastidial structures [20,47], but the exact localization of carotenoids is beyond the scope of this study and will be addressed in future research.

Several reports in the literature discuss heterologous ketocarotenoid production in plants, through nuclear or plastidial modifications, but there are no reports on the use of undifferentiated cells. Huang and colleagues expressed algal β-carotene ketolase and hydroxylase genes in tomato, resulting in ketocarotenoid accumulation (Astx = 3.12 mg g^-1^ DW) [48]. Further investigations introduced combinations of β-carotene hydroxylase and ketolase *crtZ* and *crtW* genes into various plants, including *Brassica napus* (Astx = 0.6 µg g^-1^ DW), *Solanum tuberosum* (Astx = 0.51 µg g^-1^ DW), *N. benthamiana* (Astx = 14.7 µg g^-1^ DW), and maize (0.016 mg g^-1^ DW). Soybean seeds, transformed with phytoene synthase and *crtW*/*bkt* genes, also yielded ketocarotenoid-rich transgenic seeds (Astx = 7 µg g^-1^ DW) [13,32,49–51]. Other examples are outlined in Table S3. Chloroplast genome engineering enables efficient, targeted gene insertion, and robust foreign gene expression without epigenetic changes [21,52]. Successful cases include astaxanthin production in *N. tabacum* (5.44 mg g^-1^ DW) and *Lactuca sativa* (178 µg g^-1^ fresh weight) via plastidial transformation [53,54]. Using undifferentiated cells in culture offers the advantage of high proliferation rates and the ability to adapt to various genetic and environmental modifications, making them ideal for scalable production.

### Ketocarotenoid production in BY-2 cells

Plant cell cultures have demonstrated the potential for commercial production of high-value molecules, as exemplified by the agreement between Protalix and Pfizer. The company uses carrot cell cultures to produce Elelyso®, a therapeutic enzyme indicated for the treatment of Gaucher disease [55]. Additionally, cultured carrot cells, as well as tomato and blueberry, have been shown to naturally produce pigments [56–58] but ketocarotenoid production has not been reported to date. With regard to tobacco undifferentiated cells, only other classes of pigments like anthocyanins have been produced [59]. Polturak and co-workers [30] successfully produced betalain pigments in BY-2 cells, resulting in colored *calli*. Also in BY-2 cells, Imamura and colleagues produced other betalains like betanidin, betanin and amaranthin [31].

Our strategy involved the combinatorial expression of carotenogenic genes controlled by a strong promoter. Since carotenoid production in non-photosynthetic BY-2 cells have not been previously documented, we hypothesized that enhancing the upstream components of the ketocarotenoid pathway would increase overall production of carotenoids, as shown in other studies reported. In maize and rice [17,38,60,61], the heterologous expression of phytoene synthase and desaturase genes led to increased production of lycopene and β-carotene, mediated by the plant’s endogenous lycopene β-cyclase enzyme. These enzymes, though derived from different organisms, perform identical functions as the native enzymes [62]. In BY-2 expressing the *crtI* gene, a small increase in total carotenoid content was observed, while maintaining a carotenoid profile similar to wild-type cells. Schaub and colleagues [35] also reported small amounts of β-carotene, neoxanthin and violaxanthin in transgenic BY-2 cell lines, alongside variability in gene expression between transgenic and wild-type cultures. They observed that the introduction of *crtI* did not consistently alter gene expression [35,63,64] and this variability also correlated with growth differences in the transgenic cultures [35]. The cell lines Y and YI exhibited a yellow coloration, with Y displaying a particularly strong phenotype due to substantial β-carotene accumulation. This suggests that the PSY enzyme plays a critical role in expanding the ketocarotenoid precursor pool. Cazzonelli and Pogson reported that while phytoene synthase enzyme is widely recognized as a key regulatory enzyme in the carotenoid pathway, it also represents a rate-limiting step in carotenogenesis [62].

Despite the initially low accumulation of carotenoids in BY-2 wild-type cells, the expression of the single *crtW* gene enabled the production of astaxanthin and canthaxanthin and significantly enhanced total carotenoid accumulation. This was attributed to the increased metabolic activity and up-regulation of endogenous carotenogenic metabolites. Thus, the introduction of this missing enzyme successfully completed the ketocarotenoid pathway. Transformation with *crtI* or *psy* combined with *crtW* gene produced a range of colorful cell lines, likely due to strong expression of the introduced genes. A transcriptomic analysis of selected lines will be performed to elucidate the gene expression profiles and regulatory networks underlying the observed phenotypic variations and explore the interplay between primary and secondary pathways. The IW cell lines exhibited lower levels of astaxanthin and canthaxanthin compared to the cell lines W, likely due to the diversion of the carotenoid pathway toward xanthophyll synthesis by β-carotene hydroxylase (CHYB) enzyme. Although neoxanthin and violaxanthin were not quantified, the chromatogram of the cell line IW09 (Figure S5) suggests higher levels of these xanthophylls than in line W04. Our multi-enzyme cell lines were designed to enhance product yield and regulate precursor flux. Remarkably, cell lines YIW135 and YIW141 exhibited the highest accumulation of canthaxanthin. This pigment is naturally found in low amounts, making it challenging to obtain. The production of higher levels of canthaxanthin is particularly significant in our study and the ability to generate cell lines with elevated canthaxanthin levels represents an achievement and addresses a specific interest in our research. The distinct phenotype of cell line YIW6, with lower ketocarotenoid amount, could be attributed to low *crtW* expression, enzyme kinetics, gene copy number and insertion site. A similar low ketocarotenoid accumulation was observed in *Lilium* leaves, with only 200 ng of astaxanthin per gram of fresh material [65]. Azadi and colleagues hypothesized that the low amount, like our findings in YIW6, was due to the low expression of *crtW*.

### Understanding key factors in ketocarotenoid production

The study of ketocarotenoid production within plant cell suspension cultures represents an innovative approach, involving the engineering of cell lines to facilitate the synthesis of high-value pigments. These systems ensure batch-to-batch product consistency, high containment levels, and uninterrupted production. It was not known whether undifferentiated non-photosynthetic cells had the ability to produce and store specific secondary metabolites. Our work shows that this is possible. The resulting cell lines exhibit a spectrum of distinct colors, reflecting unique combinations of carotenoid pigments. The utilization of modified tobacco BY-2 cell cultures offers a greener and economically viable alternative to chemical synthesis, as well as to extraction from natural producers, which affects the natural environment. The choice of *crtW* gene for transformation can impact ketocarotenoid production. Potential differences in efficiency between marine bacterial (CRTW) and algal (BKT) ketolases have been reported [14,21,53,66]. Other factors to consider include tissue type (e.g. roots, leaves), availability of precursors, and the expression of other enzymes that boost ketocarotenoid precursor production. Avoiding stable transformation with multiple genes can reduce the risk of gene integration failure or rearrangement, as random integration can disrupt endogenous gene function.

In microbial systems, Fraser and colleagues performed an *in vitro* characterization of marine bacterial, algae and bacterial enzymes, including β-carotene hydroxylase (CRTZ) and ketolase (CRTW and BKT). They demonstrated that marine bacterial CRTZ converted canthaxanthin to astaxanthin preferentially, whereas *Erwinia* CRTZ favored zeaxanthin formation from β-carotene. CRTW/BKT enzymes responded readily to fluctuations in substrate levels [67]. The complex conditions of marine environments, including high salinity, pressure, low temperature, and specialized lighting conditions, have prompted marine microorganisms to evolve enzyme systems that are more stable and active than those found in plants or animals. These challenges have resulted in notable distinctions between marine microbial enzymes and their terrestrial counterparts [68]. Understanding the interplay between *crtW* expression, host plant systems, and carotenoid profiles is essential, as β-carotene ketolase expression in different species had differential effects on the overall carotenoid accumulation. Variability in ketocarotenoid production may arise from transformation methods, specific genes and promoters employed, plant species, or tissue type. Coordinating these factors is crucial to attain optimal ketocarotenoid production within the selected system.

Ketocarotenoid production was previously achieved in microalgae and fermentation systems, although presenting a few drawbacks. Microalgal species frequently exhibit a slow growth rate, leading to an elevated risk of contamination during large-scale cultivation and incurring high production costs [69]. Bacteria, although they are fast-growing and economical, require engineering of the entire biosynthetic pathway, unlike our system which requires only a single gene. Yeast heterologous expression introduces challenges in protein translation and folding within a non-natural environment. Furthermore, this process relies on using metabolic intermediates produced by the host’s enzymatic machinery. In multistep pathways, efficiency may be compromised by potential losses of intermediates through diffusion, degradation, or conversion by competing enzymes [70]. Nonetheless, several successful cases have been reviewed in [71].

### Future prospects in ketocarotenoid research: paving the way for industrial-scale ketocarotenoid manufacturing

The study of ketocarotenoid pathways is relevant for understanding intrinsic cellular mechanisms, but also for applications in biotechnology. Recent advances in synthetic biology, metabolic engineering, and systems biology have opened new ways for their production, shifting from extraction from natural sources into engineered biofactories. Although microbial production is a viable option, the alternative plant cell cultures presented in this report offer a superior alternative. They have higher metabolic rates, driven by rapid cell mass proliferation and homogeneity [34,72]. This allows for the study of biosynthetic pathways and enables the formation of secondary metabolites within a short cultivation time, typically around 1-4 weeks. To the best of our knowledge, this is the first report of consistent ketocarotenoid production in non-photosynthetic undifferentiated cell cultures. Future work will validate the scalability of our system, with ongoing efforts focused on establishing pilot-scale cultures. Another emerging technology derived from BY-2 cell suspension cultures is cell-free expression systems, providing a rapid and efficient platform for producing proteins and secondary metabolites. These systems facilitate the synthesis of diverse compounds, such as lycopene, indigoidine, betanin and betaxanthins, with promising yields [73]. The versatility of this solution enables the production of valuable molecules tailored to specific industrial, pharmaceutical, or nutritional requirements, but the cost is still high. Looking ahead, the future of industrial-scale ketocarotenoid manufacturing lies in integrating AI-driven design with advanced bioreactor systems for predictive optimization and real-time production monitoring. Precision fermentation and controlled metabolic flux will improve yields, reduce costs, and address regulatory challenges. Efforts to expand substrate flexibility, such as utilizing agricultural residues, can also enhance economic feasibility and environmental sustainability of production processes.

## Experimental procedures

### Plant material

#### Tobacco BY-2 cell culture conditions

*Nicotiana tabacum* cv. Bright Yellow 2 (BY-2) cells were maintained as described in [45]. Briefly, cultures were grown in supplemented Murashige and Skoog (MS, Duchefa) medium containing 3% (w/v) of sucrose, inositol, phosphate, thiamine, and 2,4-Dichlorophenoxyacetic acid), on an orbital shaker at 28°C and 120 rpm in darkness. Stock cultures were maintained as *calli* on MS medium solidified with 0.7% agar, also kept in darkness at 28°C.

To prepare BY-2 cells for transformation with carotenogenic genes, cultures were grown under a 16-hour light (Inspire GPDL-B1-E12-09-840-01-A3PC) and 8-hour dark photoperiod, with a LED light intensity of 80-100 μmol m^-2^ s^-1^. Cell suspension cultures were renewed by transferring 3% (v/v) of the culture into 25 ml of MS medium, gradually reducing sucrose concentrations: starting at 3%, then 2%, 1%, and finally 0.5%. Simultaneously, sterilized kinetin (Duchefa) was added to the medium at concentrations ranging from 0.46 to 27.88 µM. Biomass was collected at various stages of the elicitation experiment and stored at -80°C for further analysis.

#### Growth conditions of *Nicotiana benthamiana* plants

*Nicotiana benthamiana* plants used for transient expression were grown for six weeks in a controlled chamber (Fitoclima 4600, Aralab) at 22°C, with an 8-hour light (Osram DST STICK 21W/825) and 16-hour dark photoperiod. The light intensity was maintained at 80-100 μmol m^-2^ s^-1^, and relative humidity was set to 65% to ensure optimal growth conditions.

### Vector construction

#### Phytoene synthase (*psy*) and phytoene desaturase (*crtI*)

Plasmids p326-ZmPSY1 and pHORP-PacrtI [38] (kindly offered by Changfu Zhu, Spain) were the source of the *psy* gene from maize (GenBank: AY324431.1) and the *crtI* gene from *Pantoea ananatis* (GenBank: D90087.2, *crtI* gene location: 3582-5060 bp), respectively. For construction of the plant transformation vectors, the genes were amplified by PCR using aatB1-psy and aatB2-psy primers for *psy*, and attB1-crtI and attB2-crtI primers for *crtI.* Primers used in this study are listed in Table S4. The fragments were individually subcloned into pDONR221 via Gateway recombination with BP Clonase™ II enzyme mix and then cloned into a pK2GW7 vector (kindly given by Jörg Becker, Portugal) via GATEWAY recombination with LR Clonase™ II enzyme mix (Invitrogen). This resulted in construct pY and construct pI (Figure 2) expressed under the Cauliflower Mosaic Virus 35S promoter (CaMV 35S) and kanamycin as selection marker.

#### β-carotene ketolase (*crtW*)

The coding sequence for *crtW* (GenBank: AB181388.1, *crtW*) from the marine bacterium *Brevundimonas* sp. NBRC 101024 strain SD212 was codon optimized for tobacco and synthesized by GeneCust (www.genecust.com/en/) with the addition of EcoRI and BamHI restriction sites at the 5′ and 3′ ends, respectively. It included a fused sequence encoding the plastid-transit peptide from pea (*Pisum sativum*) ribulose 1,5-biphosphate carboxylase small subunit (GenBank: X00806.1). The gene was then cloned into the pTRA vector (kindly offered by Thomas Rademacher, Aachen, Germany) using the EcoRI and BamHI sites. The resulting construct (pW) (Figure 2) contained the β-carotene ketolase gene controlled by the CaMV 35S constitutive promoter and a 35S terminator.

All three binary plasmids were transferred into *Agrobacterium tumefaciens* by the freeze-thaw method. Constructs pY and pI were inserted into strain GV3101::pMP90 while construct pW was transferred to strain GV3101::pMP90RK.

### Transient expression in *Nicotiana benthamiana* leaves

Recombinant *Agrobacterium* cells, harboring either the p19 gene sequence, the empty vector pK2GW7, or one of three constructs (pY, pI, or pW) were cultured in Yeast Extract Beef (YEB) medium. The YEB medium was composed of 0.1% (w/v) of nutrient broth, 0.1% (w/v) of yeast extract, 0.5% (w/v) of tryptone, 0.5% (w/v) of sucrose, and 2 mM of MgSO_4_, with a final pH of 7.4. The medium was supplemented with 50 mg L^-1^ of rifampicin and kanamycin, and 80 mg L^-1^ of carbenicillin for pW, while the other constructs were cultured with 50 mg L^-1^ of rifampicin and spectinomycin, and 25 mg L^-1^ of gentamycin. Cultures were incubated at 28°C and 200 rpm in the dark for a period of 2-3 days before being subcultured onto fresh medium. Four distinct bacterial suspensions were prepared by resuspending each transformed *Agrobacterium* (pY; pI; pW) in infiltration medium [74] to a final optical density at 600 nm (OD_600nm_) of 0.2. For multi-combinatorial transient expression, the transformed *Agrobacteri*a were co-cultured in infiltration medium in various combinations (pY + pI; pY + pW; pI + pW; pY + pI + pW). Each one of these bacterial suspensions was co-cultured with *Agrobacterium* harboring the p19 gene, an RNA silencing suppressor (WAK97598.1), to a final OD_600nm_ of 0.1. Fully grown *N. benthamiana* leaves were punctured on the abaxial surface, and the *Agrobacterium* suspensions were introduced using a blunt-end 1 ml syringe until the entire leaf was infiltrated. Leaves were harvested 72 h post-infection and stored at -80°C until further analysis.

### Stable transformation of tobacco BY-2 cell suspension cultures

Four-day-old tobacco BY-2 cells were transformed by co-cultivation with recombinant *A. tumefaciens*, following the protocol described by An [75], with slight modifications [76]. For multigene transformation, recombinant *A. tumefaciens* carrying individual genes were incubated together in infiltration medium for two hours before co-culturing with BY-2 cells (Table S2). After a two-day co-culture period, the cells were transferred to MS medium solidified with 0.4% gelrite (Duchefa), containing 500 mg L^-1^ ticarcillin disodium/clavulanate potassium (Timentin, Duchefa) to eliminate *Agrobacterium* and 100 mg L^-1^ kanamycin to select transformants. Cells were grown at 28°C with a 16-hour fluorescent light photoperiod and micro-*calli* were transferred to fresh medium containing 100 mg L^-1^ kanamycin and 50% decrease of Timentin after two to four weeks [76].

Following transformation, transgenic BY-2 cell cultures were grown as described above, under a 16-hour LED light photoperiod in supplemented MS medium containing 100 mg L^-1^ kanamycin (NZYTech). BY-2 *calli* were grown under identical conditions, except that the fluorescent light (Philips TL-D 36W/54-765) intensity was maintained at 50 μmol m^-2^ s^-1^ for *calli* growth. Wild-type and transgenic liquid cultures were subcultured into fresh medium weekly, while *calli* were subcultured monthly.

### Screening of positive transformants

Genomic DNA was extracted from tobacco BY-2 wild-type (WT) and transgenic cell lines with NZY Plant/Fungi gDNA Isolation kit (NZYTech, Portugal) following the manufacturer’s instructions. PCR reactions were performed using Supreme NZYTaq II 2x Colourless Master Mix (NZYTech, Portugal) with the following thermal cycling conditions: initial denaturation at 95°C for 10 min followed by 34 cycles of denaturation (94°C, 30 s), annealing (65-69°C for 30 s), extension (72°C, 15 s) followed by a final extension (72°C, 10 min). The design of primers was carried out using the Primer3 Plus software, and primer sequences and annealing temperatures are outlined in Table S4.

### Carotenoid extraction and quantification

#### Carotenoid profile analysis of BY-2 wild-type cells and *Nicotiana benthamiana leaves*

WT BY-2 cells collected during the elicitation experiment and *N. benthamiana* leaves used for transient expression were ground using liquid nitrogen. Pigments were extracted from the cells or leaves with ethanol (1:30, m/v) at 45°C for 2 h, followed by six rounds of vortexing. The mixture was centrifuged, the supernatant was collected, and the extraction repeated until color exhaustion. The organic phase was filtered through a 0.22 µm nylon syringe filter [77]. High-performance liquid chromatography (HPLC) analysis of the organic phase was carried out using a Waters Alliance System equipped with a diode array detector, with a detection range of 190 to 800 nm. A C18 reverse phase column (DeltaPaK 5µm particle size, 3.9 x 150 mm) was used for pigment separation. The mobile phase was a linear gradient of acetonitrile (ACN) and water with a flow rate of 0.5 mL min^-1^ (starting with 80% ACN for 1 min, increasing from 80% to 100% ACN over 8 min, maintained at 100% for 3 min, reduced to 80% over 2 min and maintained at 80% ACN for 6 min) [77]. The chromatograms were recorded at 445 nm and the carotenoids were identified by comparing retention time and UV absorption spectral properties to reference standards (kindly given by Hugo Pereira, Portugal) and previous analysis. Data processing was carried out using the automated integration software Empower 3.

#### Quantitative analysis of carotenoids in transgenic BY-2 cell lines

Carotenoid extraction was performed based on the methodology outlined in [78]. The cells were lyophilized for 72 hours and then ground using liquid nitrogen. A mixture of 10 mg of biomass and 500 μL of hexane:ethyl acetate (1:1) was incubated at 45°C for 2 hours, followed by six rounds of vortexing. The colored organic fraction was collected by centrifugation, and the extraction steps were repeated until color exhaustion. The organic phase was evaporated in a rotary evaporator, and the dry extract was resuspended in methanol to a final concentration of 5 mg mL^-1^. The samples were then filtered as previously described. The HPLC analysis of the collected supernatant was conducted following established protocols with slight modifications. The mobile phase was a linear gradient of acetonitrile and water, with a flow rate of 0.5 mL min^-1^ (starting at 70% ACN, increasing to 100% in 20 min, maintained at 100% for 5 min, then reduced back to 70% ACN over 2 min, and held at 70% ACN for 10 min). The chromatograms were recorded at 445 nm and, carotenoids were identified based on retention time and UV absorption spectral properties of reference standards for astaxanthin (Sigma-Aldrich), canthaxanthin (Sigma-Aldrich), and β-carotene (Sigma-Aldrich) [79]. Quantitative analysis was carried out by integrating peak areas from the chromatogram. One-way analysis of variance (ANOVA) Uncorrected Fisher’s Least Significant Difference (LSD) was used to determine significant differences between pairwise comparisons among the transgenic lines and their controls, using GraphPad Prism software (GraphPad Software).

## Authorship

All authors conceived the idea, BAR executed the experimental procedures. RA supervised the plant cell culture transformation and analysis, MRV supervised the pigment analysis. BAR wrote the initial draft with input from all authors. All authors agreed with contributed to the final version of the manuscript.

## Acknowledgments

We are grateful to the late Dr. Changfu Zhu for kindly providing the carotenogenic genes essential for this study. We also thank Dr. Jörg Becker and Dr. Thomas Rademacher for the vector backbones. Furthermore, we are grateful to several colleagues, namely André Folgado, Pedro Carvalho, Gonçalo Elias da Silva, Pedro Barros, and Inês Chaves, for providing reagents and engaging in fruitful discussions throughout the course of this work.

This work was funded by InnOValley Proof of Concept Fund (IOVPoC-2021-09) awarded by the Municipality of Oeiras. The authors are funded by Fundação para a Ciência e a Tecnologia (FCT, Portugal) through Research Units “Bioresources 4 Sustainability” (GREEN-IT, UIDB/04551/2020) and “Molecular, Structural and Cellular Microbiology” (MOSTMICRO, UIDB/04612/2020), and Plants for Life PhD Program (Fellowship PD/BD/149196/2019).

