## Supplementary Information for "Metabolically engineered plant cell cultures as biofactories for the production of high-value carotenoid pigments astaxanthin and canthaxanthin"

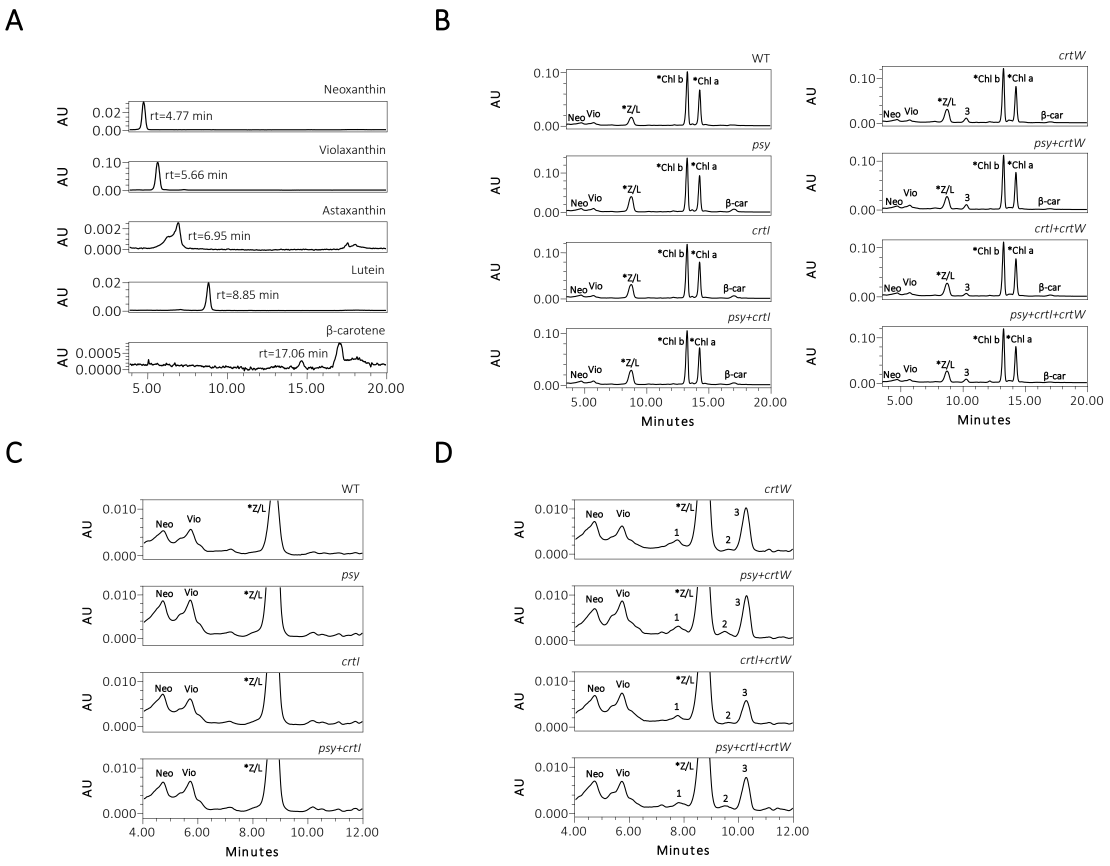


Figure S1. Carotenoid profile of *N. benthamiana* Agro-infiltrated leaves collected at 3 days post-infiltration. **(A-B)** Two to three leaves were Agro-infiltrated with each construct. Peaks were identified as neoxanthin (Neo), violaxanthin (Vio) and β-carotene (β-car) by comparing retention times and overlaying chromatograms with those of corresponding standards **(A)**. Additionally, putative lutein/zeaxanthin (Z/L*), chlorophyll a (*Chl a) and chlorophyll b (*Chl a, *Chl b) were identified by comparing chromatographic profiles **(B)** to analyzed carotenoid profiles from Arabidopsis and *Tetraselmis chuii*, as well as descriptions in the literature. In the chromatograms close-up comparing xanthophyll- **(C)** and ketocarotenoid-producing leaves **(D)**, unidentified peaks 1, 2 and 3 are visible.

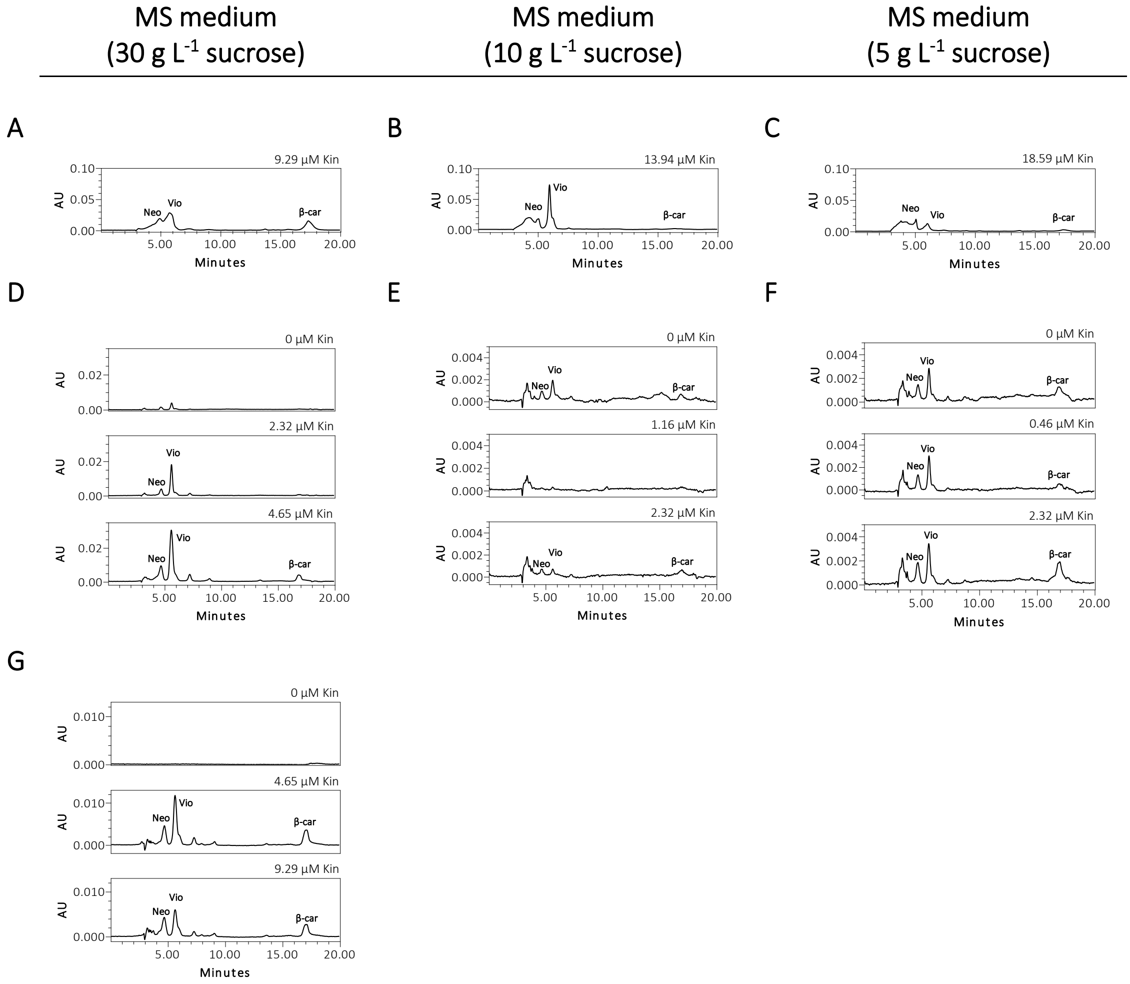


**Figure S2.** Carotenoid production in tobacco BY-2 WT cell suspension cultures. HPLC analysis of the carotenoid profile of cells grown under a 16 h-light photoperiod for **(A-C)** 4 months, **(D-F)** 10 months, and **(G)** 14 months. Cells were subcultured in MS medium formulated with varying amounts of sucrose concentrations (3%, 1% and 0.5%) and kinetin (Kin, 0.46 – 18.59 µM) until the 10-month time point. After this analysis, we selected cultures grown with 3% of sucrose, which showed higher yields in total carotenoid content. Sample injection volumes for HPLC were **(A-C)** 100 µL, **(D-F)** 20 µL and **(G)** 10 µL.

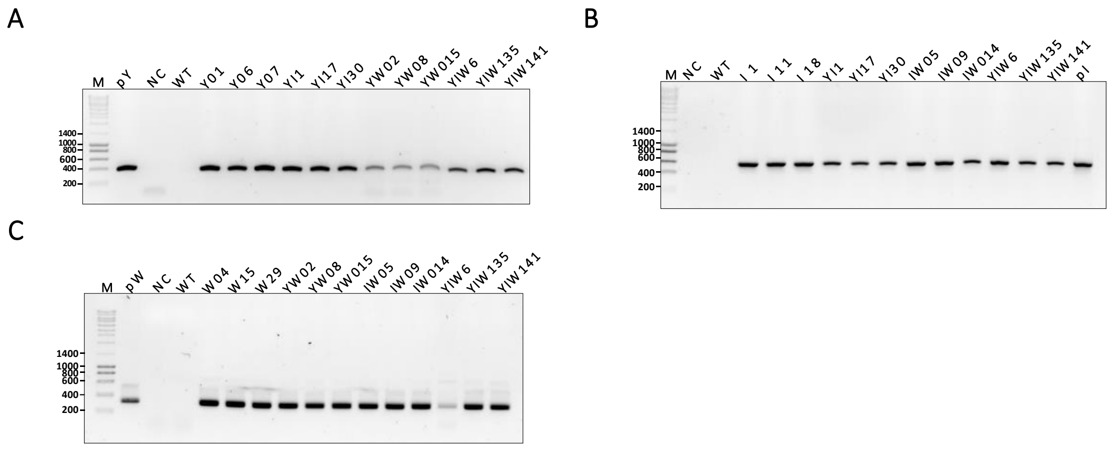


**Figure S3.** PCR analysis of genomic DNA of BY-2 cell lines. PCR amplified product of **(A)** maize phytoene synthase (*psy*) (408 bp), **(B)** bacterial phytoene desaturase (*crtI*) (579 bp) and **(C)** bacterial β-carotene ketolase (*crtW*) (331 bp) genes. M: molecular size marker; NC: Negative control; pY, pI, pW: construct pY, construct pI, construct pW, respectively; WT: wild-type.

A

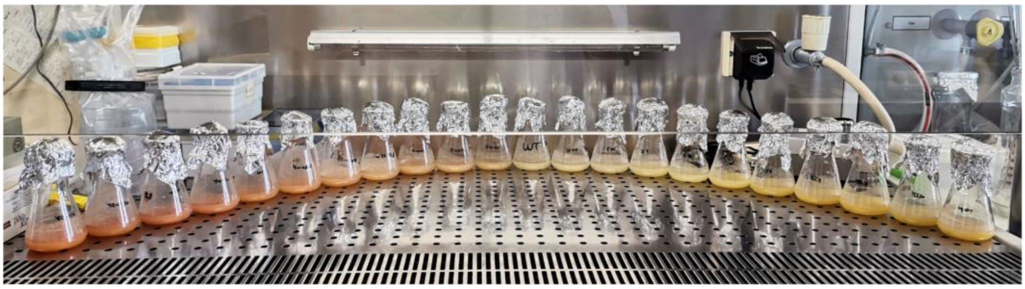

B

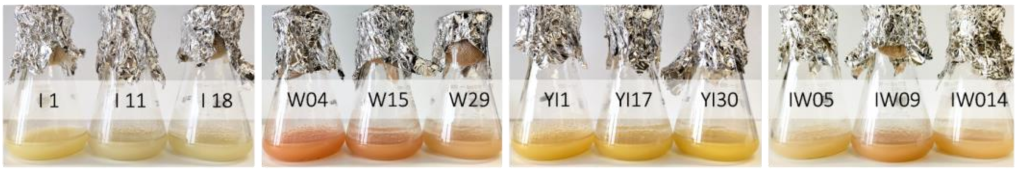


C

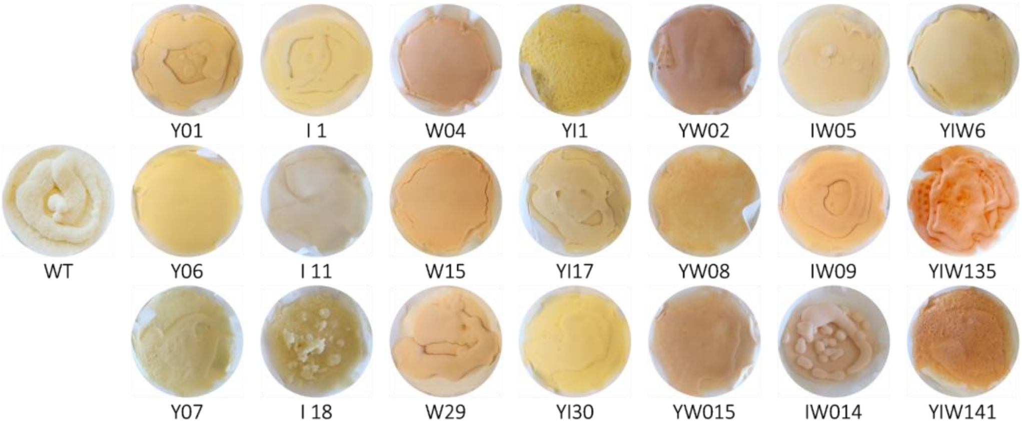


**Figure S4**. Tobacco BY-2 cultured cells producing ketocarotenoids and/or xanthophylls. **(A)** Cell cultures are shown according to the color observed by visual inspection, ranging from orange (left) to salmon, pale WT (center), light yellow to dark yellow (right). **(B)** Cell suspension cultures and **(C)** filtered liquid cultures used in this study.

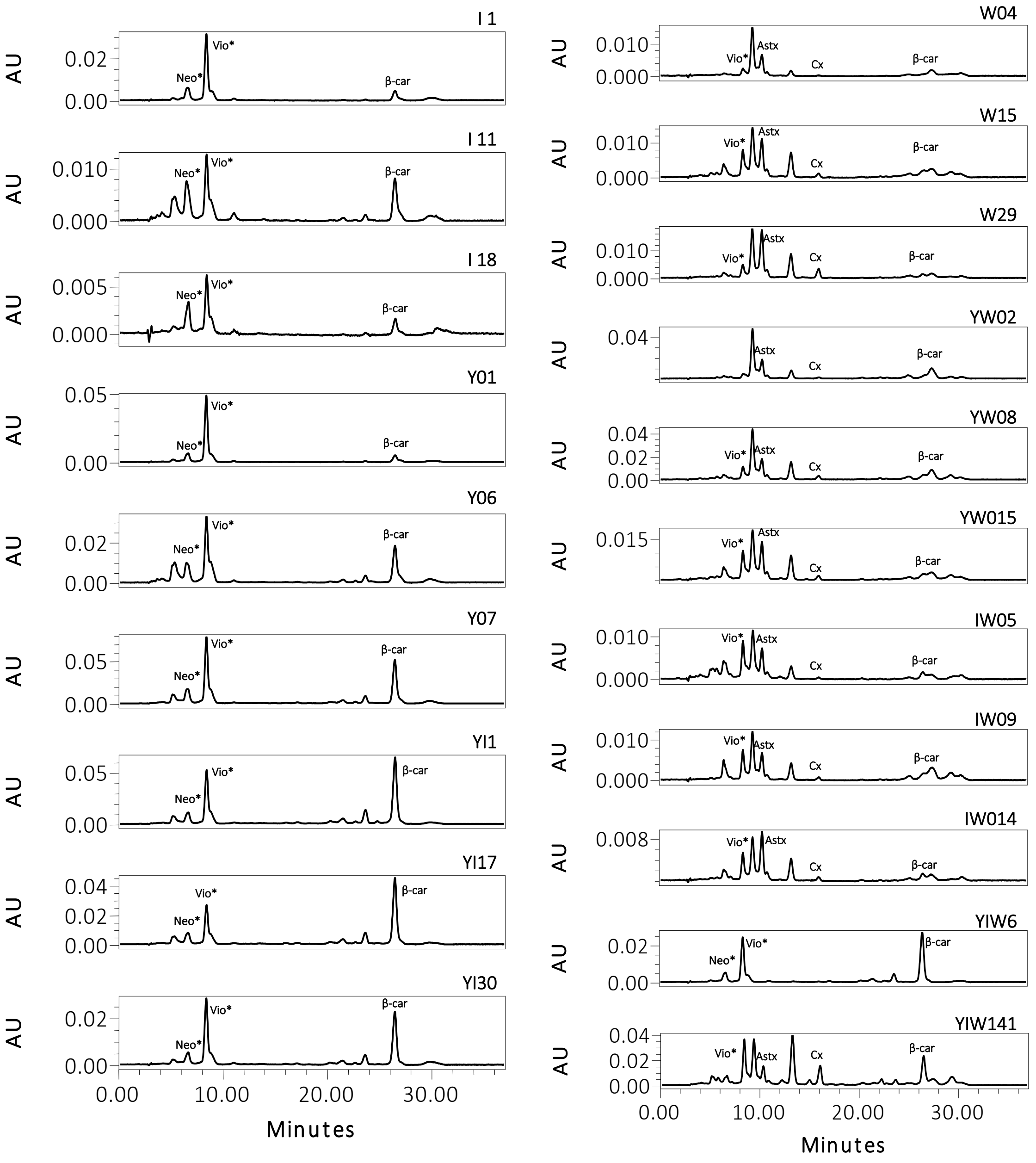


**Figure S5.** Carotenoid profile of tobacco BY-2 transgenic cell lines. The chromatograms were recorded at 445 nm, and the y-axis was scaled to the highest peak for clarity. Carotenoids were identified as follows: putative neoxanthin (*Neo), putative violaxanthin (*Vio), astaxanthin (Astx), canthaxanthin (Cx) and β-carotene (β-car).

**Table S1.** Overview of the experimental conditions employed for the carotenoid elicitation experiment in tobacco BY-2 cells.

| | Sucrose (g $L^{-1}$) | Kinetin (µM) | Inoculum (v/v) |
|---|---|---|---|
| Treatment in the dark | 30 | 0 | 3% |
| | **Sucrose (g $L^{-1}$)** | **Kinetin (µM)** | **Inoculum (v/v)** |
| Treatment in the light | 30 | 0 | 3% |
| | 30 | 4.65 – 27.88 | |
| | 10 | 0 | 3-8% |
| | 10 | 1.16 – 13.94 | |
| | 5 | 0 | 3-8% |
| | 5 | 2.32 – 18.59 | |

**Table S2.** Summary of the main parameters of Agrobacterium-mediated transformation of tobacco BY-2 cells.

| Genotype | *A. tumefaciens* strain (binary vector) | Co-culture* $OD_{600} = 0.5$ | Selected cell lines |
|---|---|---|---|
| Y | GV3101::pMP90 (pK2GW7:ZmPSY1 (pY)) | No | Y01, Y06, Y07 |
| I | GV3101::pMP90 (pK2GW7:PacrtI (pI)) | No | I 1, I 11, I 18 |
| W | GV3101::pMP90RK (pTRA:crtW (pW)) | No | W04, W15, W29 |
| YI | GV3101::pMP90 (pK2GW7:ZmPSY1 (pY))<br>GV3101::pMP90 (pK2GW7:PacrtI (pI)) | Yes | YI1, YI17, YI30 |
| YW | GV3101::pMP90 (pK2GW7:ZmPSY1 (pY))<br>GV3101::pMP90RK (pTRA:crtW (pW)) | Yes | YW02, YW08, YW015 |
| IW | GV3101::pMP90 (pK2GW7:PacrtI (pI))<br>GV3101::pMP90RK (pTRA:crtW (pW)) | Yes | IW05, IW09, IW014 |
| YIW | GV3101::pMP90 (pK2GW7:ZmPSY1 (pY))<br>GV3101::pMP90 (pK2GW7:PacrtI (pI))<br>GV3101::pMP90RK (pTRA:crtW (pW)) | Yes | YIW6, YIW135, YIW141 |

* Agrobacteria cultures were individually collected by centrifugation and subsequently resuspended together in a solution of infiltration medium, which was further supplemented with 200 µM acetosyringone.

**Table S3.** Representative list of heterologous production of astaxanthin and canthaxanthin, highlighting the highest yields reported.

| Species | Ketolase gene | Origin | Astaxanthin | Canthaxanthin | Tissue | Reported source |
|---|---|---|---|---|---|---|
| *Arabidopsis thaliana* | *bkt*[a] | *Chlamydomonas reinhardtii, Chlorella zofingiensis, Haematoccocus pluvialis* | 12 – 2070 µg g$^{-1}$ DW | 200 – 600 µg g$^{-1}$ DW | Leaf, seed | [1] |
| *Brassica napus* | *crbkt*[b] | *C. reinhardtii* | 28.2 – 33.2 µg g$^{-1}$ FW | 4 – 4.3 µg g$^{-1}$ FW | Cotyledon petioles | [2] |
| *Daucus carota* | *bkt*[b] | *H. pluvialis* | 12.4 – 91.6 µg g$^{-1}$ FW | 4 – 50.1 µg g$^{-1}$ FW | Callus, leaf, root | [3] |
| *Glycine max* | *bkt*[b], *crtW*[b] | *Brevundimonas* sp. SD212, *H. pluvialis* | 2 – 7 µg g$^{-1}$ DW | 4 – 52 µg g$^{-1}$ DW | Seed | [4] |
| *Lotus japonicus* | *crtW* | *Agrobacterium aurantiacum* | 89.8 µg g$^{-1}$ FW | | Flower | [5] |
| *Lactuca sativa* | *crtW*[b] | *Brevundimonas* sp. SD212 | 178 µg g$^{-1}$ FW | 12 µg g$^{-1}$ FW | Leaf | [6] |
| *Lilium x formolongi* | *crtW*[b] | *Brevundimonas* sp. SD212 | 0.2 – 0.5 µg g$^{-1}$ FW | 0.6 – 3.5 µg g$^{-1}$ FW | Callus, leaf | [7] |
| *Nicotiana benthamiana* | *CBFD, HBFD* | *Adonis aestivalis* | 0.15 – 0.99 mg g$^{-1}$ DW | 0.003 mg g$^{-1}$ DW | Leaf | [8] |
| | *crtW* | *Brevundimonas* sp. SD212 | 0.13 – 0.4 mg g$^{-1}$ DW | 0.003 – 0.012 mg g$^{-1}$ DW | | |
| | *crtW*[b] | *Brevundimonas* sp. SD212 | 3.6 –14.7 µg g$^{-1}$ DW | 2.9 – 162.3 µg g$^{-1}$ DW | Leaf | [9] |

| Species | Ketolase gene | Origin | Astaxanthin | Canthaxanthin | Tissue | Reported source |
|---|---|---|---|---|---|---|
| *Nicotiana glauca* | *crtO*[a], *crtW* | *Synechocystis, Nostoc* 73102 | - | - | Flower | [10] |
| | *crtW*[b] | *Brevundimonas* sp. SD212 | 0.04 – 0.29 µg mg$^{-1}$ DW | 0.03 – 1.05 µg mg$^{-1}$ DW | Leaf, ovary, petal | [11] |
| *Nicotiana tabacum* | *crtO*[a] | *H. pluvialis* | 2.4 – 83.9 µg g$^{-1}$ FW | 36.4 mg g$^{-1}$ FW | Nectary, leaf | [12] |
| | *crtW*[b] | *Brevundimonas* sp. SD212 | 1.88 – 5.44 mg g$^{-1}$ DW | 0.08 – 0.14 mg g$^{-1}$ DW | Leaf | [13] |
| | *crtW*[b] | *Paracoccus sp.* | 0.064 – 0.8 mg g$^{-1}$ DW | | Leaf, nectary | [14] |
| *Solanum lycopersicum* | *crtW*[b] | *Brevundimonas* sp. SD212 | 47 – 83 µg g$^{-1}$ DW | 8 – 899 µg g$^{-1}$ DW | Fruit | [15] |
| | | | 38.7 a 362.5 µg g$^{-1}$ DW | 19.8 – 887 µg g$^{-1}$ DW | Fruit, leaf | [16] |
| | *bkt*[b] | *C. reinhardtii* | 25.2 – 92.4 µg g$^{-1}$ DW | 26.1 – 84.6 µg g$^{-1}$ DW | Fruit | [17] |
| | | | 0.35 – 3.12 mg g$^{-1}$ DW | 1.59 – 2.3 mg g$^{-1}$ DW | Fruit, leaf | [18] |
| | *crtW*[a,b] | *Brevundimonas* sp. | 0.2 mg g$^{-1}$ DW | 1.5 mg g$^{-1}$ DW | Fruit | [19] |
| *Solanum tuberosum* | *crtO* | *Synechocystis sp.* | 0.7 – 1.8 µg g$^{-1}$ DW | - | Tuber | [20] |
| | *bkt* | *H. pluvialis* | 0.2 – 13.9 µg g$^{-1}$ DW | 0.2 µg g$^{-1}$ DW | Tuber | [21] |
| | *crtW*[b] | *Brevundimonas* sp. SD212 | 0.3 – 32.3 µg mg$^{-1}$ DW | 0.06 – 0.09 µg mg$^{-1}$ DW | Leaf, tuber | [22] |

| Species | Ketolase gene | Origin | Astaxanthin | Canthaxanthin | Tissue | Reported source |
|---|---|---|---|---|---|---|
| *Oryza sativa* | *bkt*[b] | *C. reinhardtii* | 0.2 – 16.2 μg $g^{-1}$ DW | 0.04 – 25.8 μg $g^{-1}$ DW | Grain | [23] |
| *Zea mays* | *bkt*[b] | *C. reinhardtii* | 10.8 – 16.8 μg $g^{-1}$ DW | | Endosperm | [24] |
| *Chlamydomonas reinhardtii* | *bkt*[b] | *cis* | 0.8 – 23.5 mg $L^{-1}$ | 0.8 – 5.2 mg $L^{-1}$ | | [25] |
| *Cyanidioschyzon merolae* | *bkt*[b] | *C. reinhardtii* | 0.45 – 0.69 wt % | 0.39 – 0.60 wt %, | | [26] |
| *Saccharomyces cerevisiae* | *bkt, crtW* | *Brevundimonas vesicularis, Bradyrhizobium sp., H. pluvialis* | 20 – 70 μg $L^{-1}$ | 69 – 1068 μg $L^{-1}$ | | [27] |
| | *bkt*[b] | *H. pluvialis* | 446.4 mg $L^{-1}$ | 1 – 5 mg $g^{-1}$ DW | | [28] |
| | | | 0.5 – 4.7 mg $g^{-1}$ DW | - | | [29] |
| *Yarrowia lipolytica* | *cbfd, hbfd*[b] | *A. aestivalis* | 0.4 – 3.5 mg $L^{-1}$ | - | | [30] |
| | *crtW*[b] | *Paracoccus* sp., | 10 – 858 mg $L^{-1}$ | - | | [31] |
| | *crtW* | *C. reinhardtii, H. pluvialis* | 0.15 – 41.3 mg $g^{-1}$ DW | 1 – 4 mg $g^{-1}$ DW | | [32] |
| | *cbfd, hbfd*[b] | *A. aestivalis* | trace amounts | | | |
| *Corynebacterium glutamicum* | *crtW*[a,b] | *Fulvimarina pelagi* | 25 – 103 mg $L^{-1}$ | 10 – 75 mg $L^{-1}$ | | [33] |

| Species | Ketolase gene | Origin | Astaxanthin | Canthaxanthin | Tissue | Reported source |
|---|---|---|---|---|---|---|
| *Escherichia coli* | *crtW*[b] | *Brevundimonas* sp. SD212 | 10 – 1180 mg $L^{-1}$ | 170 – 1100 mg $L^{-1}$ | | [34] |
| | *crtW*[b] | *Brevundimonas* sp. SD212 | 75.6 – 610.4 μg $g^{-1}$ DW | 44.7 – 92 μg $g^{-1}$ DW | | [9] |
| | *crtW*[a,b] | *Anabaena variabilis* *Brevundimonas* sp. SD212 | 70 – 320 mg $L^{-1}$ | 1 – 10.1mg $L^{-1}$ | | [35] |
| *Mucor circinelloides* | *bkt*[b] | *H. pluvialis* | 2 – 41 μg $g^{-1}$ DW | 63 – 576 μg $g^{-1}$ DW | | [36] |
| Human embryonic kidney cells (HEK293T) | *crtW*[b], *bkt*[b] | *Brevundimonas* sp. *C. reinhardtii* *Haematococcus lacustris* | 3.1 – 232.8 μg $g^{-1}$ DW | 29.3 – 153.1 μg $g^{-1}$ DW | | [37] |

[a] A strain or cultivar that has been previously modified; [b] Other genes were employed in a multi-transformation approach; DW: dry weight, FW: fresh weight.

Table S4. Primers used in the present study for vector construction and amplification of phytoene synthase, phytoene desaturase and β-carotene ketolase gene fragments.

| Primer name | Nucleotide sequence | G+C content (%) | $T_{ann}$ (°C) / Elongation time (s) |
|---|---|---|---|
| attB1-psy | 5'-GGG GAC AAG TTT GTA CAA AAA AGC AGG CTT CAT GGC CAT CAT ACT CGT AC-3' | 46 | 60 / 90 |
| attB2-psy | 5'-GGG GAC CAC TTT GTA CAA GAA AGC TGG GTC CTA GGT CTG GCC ATT TCT CA-3' | 52 | |
| attB1-crtI | 5'-GGG GAC AAG TTT GTA CAA AAA AGC AGG CTG CAT GGC TTC TAT GAT ATC CT-3' | 44 | 56 / 90 |
| attB2-crtI | 5'-GGG GAC CAC TTT GTA CAA GAA AGC TGG GTC TCA TAT CAG ATC CTC CAG CA-3' | 50 | |
| psy_F | 5'-ACA TTC AGC CAT TCA GGG ACA-3' | 48 | 65 / 15 |
| psy_R | 5'-TTA CCC CTC TCT CTG CCT CC-3' | 60 | |
| crtI_F | 5'-TAA TTG GTG CAG GCT TCG GT-3' | 50 | 69 / 30 |
| crtI_R | 5'-ATG AGG TGG CGA AGG GAT TG-3' | 55 | |
| crtW_F | 5'-GGC TGA ACC AAG AAT TGT GCC-3' | 52 | 68 / 10 |
| crtW_R | 5'-ACC TGG AGC TGC ATG ATG AG-3' | 60 | |

F Forward, R reverse

of the carotenoid biosynthesis pathway. *Plant Biotechnol. Rep.* 4, 269–280. https://doi.org/10.1007/s11816-010-0147-y

Biosynthesis in Rice Endosperm. *Mol. Plant* 11, 1440–1448. https://doi.org/10.1016/j.molp.2018.09.007
